# Detecting task-dependent modulation of spatiotemporal module via tensor decomposition: application to kinematics and EMG data for walking and running at various speed

**DOI:** 10.1101/700872

**Authors:** Ken Takiyama, Hikaru Yokoyama, Naotsugu Kaneko, Kimitaka Nakazawa

## Abstract

How the central nervous system (CNS) controls many joints and muscles is a fundamental question in motor neuroscience and related research areas. An attractive hypothesis is the module hypothesis: the CNS controls groups of joints or muscles (i.e., spatial modules) while providing time-varying motor commands (i.e., temporal modules) to the spatial modules rather than controlling each joint or muscle separately. Another fundamental question is how the CNS generates numerous repertories of movement patterns. One hypothesis is that the CNS modulates the spatial and/or temporal modules depending on the required tasks. It is thus essential to quantify the spatial module, the temporal module, and the task-dependent modulation of those modules. Although previous methods attempted to quantify these aspects, they considered the modulation in only the spatial or temporal module. These limitations were possibly due to the constraints inherent to conventional methods for quantifying the spatial and temporal modules. Here, we demonstrate the effectiveness of tensor decomposition in quantifying the spatial module, the temporal module, and the task-dependent modulation of these modules without such limitations. We further demonstrate that the tensor decomposition provides a new perspective on the task-dependent modulation of spatiotemporal modules: in switching from walking to running, the CNS modulates the peak timing in the temporal module while recruiting proximal muscles in the corresponding spatial module.

**Author summary:** There are at least two fundamental questions in motor neuroscience and related research areas: 1) how does the central nervous system (CNS) control many joints and muscles and 2) how does the CNS generate numerous repertories of movement patterns. One possible answer to question 1) is that the CNS controls groups of joints or muscles (i.e., spatial modules) while providing time-varying motor commands (i.e., temporal modules) to the spatial modules rather than controlling each joint or muscle separately. One possible answer to question 2) is that the CNS modulates the spatial and/or temporal module depending on the required tasks. It is thus essential to quantify the spatial module, the temporal module, and the task-dependent modulation of those modules. Here, we demonstrate the effectiveness of tensor decomposition in quantifying the modules and those task-dependent modulations while overcoming the shortcomings inherent to previous methods. We further show that the tensor decomposition provides a new perspective on how the CNS switches between walking and running. The CNS modulated the peak timing in the temporal module while recruiting proximal muscles in the corresponding spatial module.

## 1 Introduction

How the central nervous system (CNS) controls our body is a fundamental question. The human body possesses 360 joints and more than 650 muscles; the CNS controls our body while somehow resolving a significant number of degrees of freedom (DoFs) [1]. For example, in walking and running in daily life, these large numbers of joints and muscles should be controlled in orchestrated manners. The module hypothesis is an influential hypothesis for how to manage the tremendous number of DoFs [2–6]. In the hypothesis, the CNS reduces the DoFs while controlling groups of joints or muscles, referred to as spatial modules, rather than single joints or muscles separately. Accordingly, the recruitment pattern of each spatial module that changes over time is referred to as a temporal module.

How the brain achieves various repertoires of motions is another fundamental question. One possible solution is that the brain modulates the spatiotemporal modules depending on the task at hand. The temporal module rather than the spatial module can be modulated depending on the task such as walking, running, various types of gaits, and responding to unpredictable perturbations while walking [7, 8]. On the other hand, the spatial module rather than the temporal module can achieve the task-dependent modulation in responding to unpredictable perturbations to maintain standing balance [9] and in arm-reaching movements [10] as well as in various types of movements of frogs [11]. In our recent finding, the number of modules showed task-dependent modulation in walking and running at various speeds [12]. Summarizing, there are several seemingly different perspectives on how the spatiotemporal modules are modulated depending on the task.

One possible reason why there is no agreed-upon perspective on the task-dependent modulations of the spatiotemporal modules is the limitation inherent to conventional methods. Conventional methods, such as principal component analysis (PCA, [13]) and non-negative matrix factorization (NNMF, [14]), are classified as matrix decomposition methods. Although matrix decomposition is suitable for discussing two factors inherent to joint angle and electromyographic (EMG) data (i.e., spatial and temporal modules), there can be limitations to considering more than three factors such as spatial modules, temporal modules, and the task-dependent modulations of those modules. For example, with matrix decomposition, the task-dependent modulation of the temporal module between two tasks has been discussed under the constraint of the same spatial module; on the other hand, the modulation of the spatial module has been discussed without considering the temporal module. These constraints can thus provide diverse and non-unified perspectives on the task-dependent modulation of the spatiotemporal modules.

Here, we demonstrate the effectiveness of tensor decomposition in extracting the spatial module, the temporal module, and the task-dependent modulations of the spatiotemporal modules. Tensor decomposition is a generalized version of matrix decomposition. By constructing tensor data that combine matrix data in the third dimension, tensor decomposition enables the extraction of three factors. In contrast to matrix decomposition, tensor decomposition thus enables the extraction of the task-dependent modulations of the spatiotemporal modules while simultaneously extracting the spatial and temporal modules. Notably, tensor decomposition can be extended to data with more than three dimensions and can extract more than three components.

Throughout this study, we rely on CANDECOMP/PARAFAC decomposition, or CP decomposition [15, 16], because it is a natural extension of conventional methods (i.e., PCA and NNMF). Concretely, CP decomposition provides spatial and temporal modules in a similar manner to conventional methods; additionally, CP decomposition generates task-dependent modulations of these modules. Another important aspect is that CP decomposition does not require orthogonality among spatial modules, in contrast to PCA. Because the orthogonality in PCA is for mathematical convenience rather than a requirement for the analysis of joint angles, CP decomposition can generate more plausible spatial modules than PCA. Furthermore, CP decomposition is applicable to non-negative data (e.g., EMG data) with non-negative constraints. Based on the above-mentioned reasons, we utilized CP decomposition, a tensor decomposition algorithm, to extract the spatial modules, the temporal modules, and the task-dependent modulations of the spatiotemporal modules.

A few studies have attempted to demonstrate the effectiveness of tensor decomposition in analyzing the joint angle and EMG data [17–19]. A previous study applied tensor decomposition to wrist EMG data and demonstrated the task-dependent modulation of the spatiotemporal module [19]. Despite the detailed description of mathematical and historical aspects of tensor decomposition, a description of movements and tasks is lacking; the significance of the tensor decomposition in discussing the task-dependent modulation of the spatiotemporal module is unclear. Other studies [17,18] applied a matrix tri-factorization to EMG data and revealed how each spatiotemporal module was modulated depending on the task. Although the tri-factorization algorithm and their new experimental paradigm [18] are sophisticated, the algorithm is a reduced version of tensor decomposition (a detailed description is given in [19]). Additionally, although the modulation patterns were estimated in an unsupervised manner (i.e., no experimental information is necessary), additional analysis for interpreting the modulation was based on a supervised method (i.e., experimental information should be required) [17,18]. Although one advantage of tensor decomposition is its feature of unsupervised learning, the requirement of supervised learning reduces the advantage provided by this method. In summary, the effectiveness of tensor decomposition is still unclear.

We apply tensor decomposition to joint angle and EMG data in walking and running at various speeds to elucidate upon how spatiotemporal modules are modulated depending on both speed and either walking or running. From the perspective of joint angle data, joint angles exhibit a speed dependence, and this dependence differs for each joint angle [3, 20]. One interesting perspective is that the spatial module is confined to a low-dimensional space independent of speed [3]. A later study suggested that the number of spatial modules depends on the speed [21]; the number at higher speeds is less than that at lower speed, and this tendency is diminished in Parkinson’s disease. From the perspective of EMG data, muscle activities show a speed dependence, and the dependence differs in each muscle [20]. With the same spatial modules, the temporal modules were suggested to be modulated depending on speed [4, 22]. In particular, a peak of some temporal modules can exhibit temporal shifts depending on whether either walking or running is occurring [22]. The temporal module and activities of certain individual muscles have been shown to differ depending on speed [4, 22], similar to the results of another study [20]. In summary, there are various perspectives on how the joint angles and EMG are modulated depending on speed, with no consensus yet to be achieved.

We demonstrate that the tensor decomposition enables us to clarify the task-dependent modulations of spatiotemporal modules inherent in joint angles and EMG data; in particular, we demonstrate the effectiveness in the analysis of walking and running at various speeds. In joint angles, we extract two types of modules: 1) the more significant recruitment at higher speeds and 2) recruitment mainly during running. From the EMG data, we demonstrate three types of modules: 1) the more significant recruitment at higher speeds, 2) recruitment mostly during walking, and 3) recruitment mainly during running. By comparing the second and third types of modules inherent to EMG data, we provide a new perspective on how the spatiotemporal modules depend on either walking or running: the CNS switches between walking and running by not only modulating the temporal module, as mentioned in previous studies [4, 22], but also recruiting proximal muscles more so during running than during walking.

## 2 Results

### 2.1 Tensor decomposition

We applied tensor decomposition to investigate the task-dependent modulation of the spatiotemporal module inherent to the joint angle and EMG data. The differences between the tensor decomposition and the matrix decomposition are in how one makes the original data able to be analyzed and the obtained results. For the tensor decomposition, the original data consist of a 3-dimensional array in the current study. Notably, an array with greater than 3 dimensions is available for tensor decomposition. The array consists of the joint or muscle sequence (*S* columns in Fig. 1a), the temporal series (*T* rows in Fig. 1a), and the task sequence (*K* slices of the *S* × *T* matrices in Fig. 1a). Throughout this study, the word “task” broadly indicates motion under all types of conditions (i.e., walking or running at different speeds) for all subjects. The tensor decomposition enabled the extraction of not only spatial and temporal modules but also task-dependent modulations of those modules (Fig. 1a). Throughout this study, we use bar graphs to denote spatial modules, line plots to denote temporal modules, and circular dots to indicate the task-dependent modulations, as denoted in Fig. 1, following a previous study [16].

**Fig. 1.**
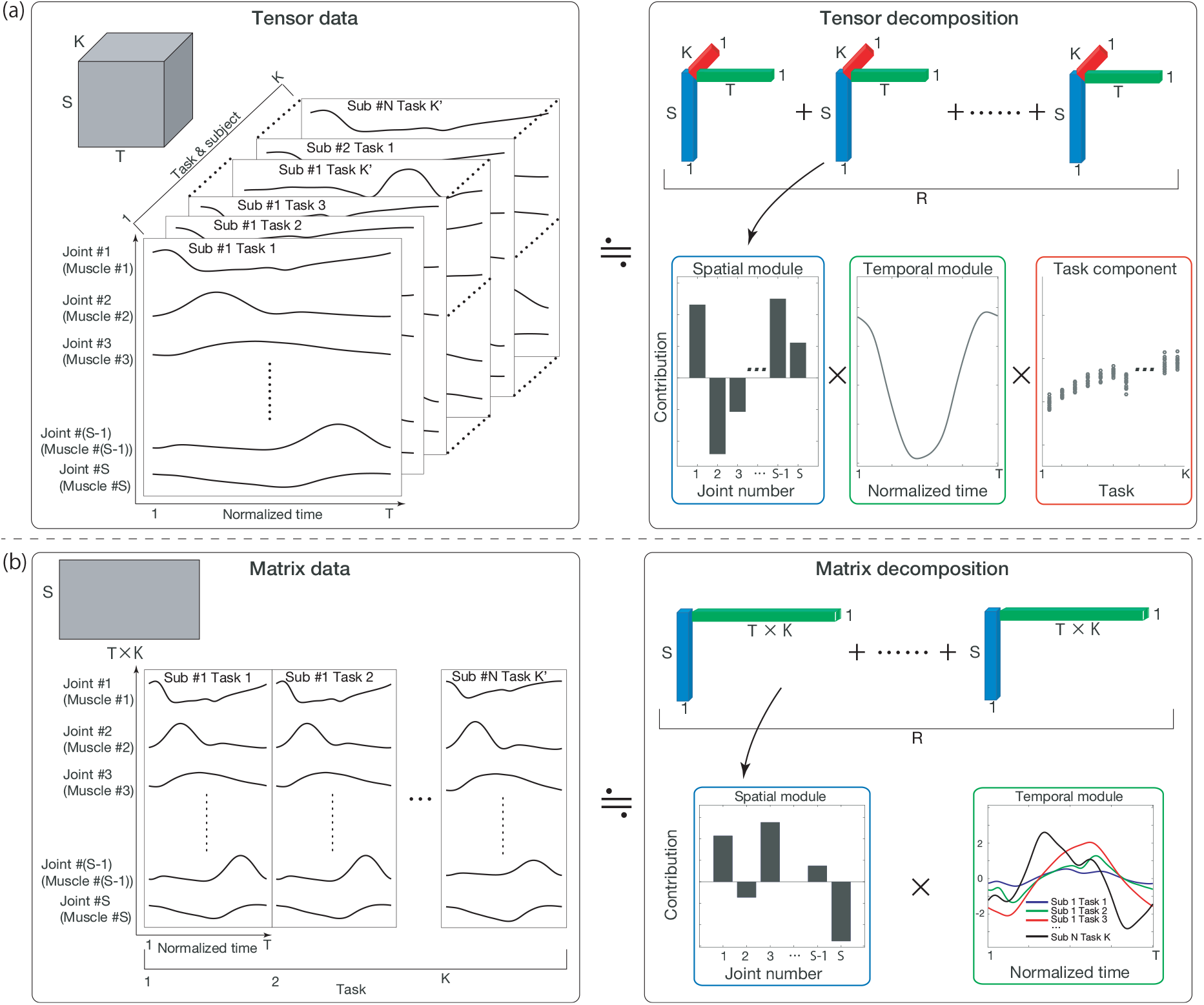
The tensor and matrix decomposition concepts. (a): The tensor decomposition and CP decomposition that we focused on in the current study. In the decomposition, we make the 3-dimensional array data consist of *S* columns, *T* rows, and *K* slices. After decomposition, we obtain the spatial module shown as a bar graph in the blue frame, the temporal module shown as a line plot in the green frame, and the task-dependent modulations of those modules shown as circle dots in the red frame. (b): Matrix decomposition. In analyzing *K* task datasets at once, we need to establish an *S* × (*T* × *K*) matrix. After applying the matrix decomposition, we obtain the spatial module and the task-dependent modulations of the temporal modules.

We focus on CP decomposition throughout this paper. In CP decomposition, the (*i, j, k*)th element of the tensor data ***X*** ∈ **R**^*S*×*T*×*K*^, *X*_*i,j,k*_, is approximated as

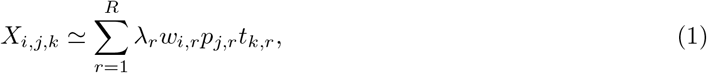

where *S*, *T*, and *K* indicate the number of joint angles or muscles, the number of time frames, and the number of tasks, respectively; *R* is the number of modules and components to be determined *a priori*; *w*_*i,r*_ indicates the *i*th element of the *r*th spatial module ***w***_*r*_ ∈ **R**^*S*×1^; *p*_*j,r*_ indicates the *j*th element of the *r*th temporal module ***p***_*r*_ ∈ **R**^*T* × 1^; *t*_*k,r*_ indicates the *k*th element of the *r*th task component ***t***_*r*_ ∈ **R**^*K* × 1^ (Fig. 1a); and *λ*_*r*_ indicates the scaling factor for the *r*th component under the conditions 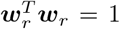, 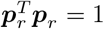, and 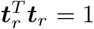. λ_*r*_ indicates the contribution of the *r*th tensor to explain original tensor data. Data at the *k*th task can be approximated as

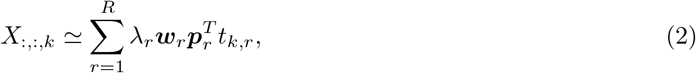

which indicates that spatiotemporal modules are common across all the tasks and that the recruitment patterns of those modules are modulated depending on *t*_*k,r*_. The spatial modules, temporal modules, and task components are estimated to minimize the squared error between the original tensor data and the decomposed data

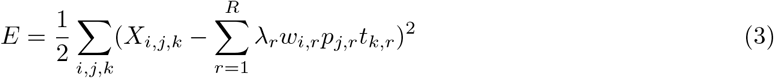

with some constraints on λ_*r*_ ≥ 0, *w*_*i,r*_, *p*_*j,r*_, and *t*_*k,r*_. There are no constraints for the analysis of joint angle and non-negative constraints for the analysis of the EMG data (i.e., *w*_*i,r*_ ≥ 0, *p*_*j,r*_ ≥ 0, and *t*_*k,r*_ ≥ 0). Throughout this study, we determined *R* as the minimum number of modules and components that explained more than 70% of the variance of the original data. Although we relied on the variance as a measure to determine *R*, following previous studies using matrix decomposition, we also demonstrated a common measure to determine *R* in tensor decomposition, with the fitting error defined as 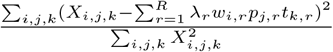.

In contrast to tensor decomposition, the matrix decomposition (e.g., PCA or NNMF) enabled the extraction of the spatial module and task-dependent modulation of only the temporal module when we analyzed *S* ×(*T* × *K*) matrices (Fig. 1b). In the decomposition, the matrix ***Z*** ∈ **R**^*S* ×(*T* × *K*)^ is decomposed as

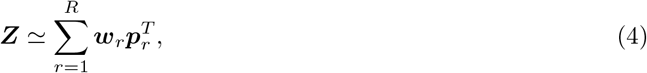

in which *S*, *T*, and *K* indicate the number of joint angles or muscles, the number of time frames, and the number of tasks, respectively; *R* is the rank to be determined *a priori*; ***w***_*r*_ ∈ **R**^*S* × 1^ indicates the *r*th spatial module; and ***p***_*r*_ ∈ **R**^(*T* × *K*)×1^ indicates the *r*th temporal module modulated in a task-dependent manner (Fig. 1b). Similar to tensor decomposition, we determined *R* as the minimum number of modules and components that explained more than 70% of the variance of the original data. In PCA, there are orthogonality restrictions among spatial modules: 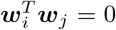 when i ≠ *j*. In NNMF, there are no such orthogonality restrictions with non-negative constraints (i.e., *w*_*i,r*_ > 0 and *p*_*j,r*_ > 0). After the analysis, we were able to consider how to evaluate the task-dependent modulations of the temporal modules (Fig. 1b). It is possible to apply the matrix decomposition to the *S* × *T* matrices in each task. In that case, we would obtain different spatial and temporal modules in each task, resulting in the fact that we should consider how to evaluate the task-dependent modulation of the spatiotemporal modules. One way is to assess the correlation of spatial modules among the tasks without evaluating the task-dependent modulation of the temporal modules. In the matrix decomposition, we need to evaluate the task-dependent modulation of either the spatial module or the temporal module.

In summary, the tensor decomposition enables the evaluation of the task-dependent modulations of the spatiotemporal modules without being restricted to considering only spatial or temporal modules.

### 2.2 Tensor decomposition for joint angle data

The current study focused on the hip, knee, and ankle angles of the right and left legs in the sagittal plane, listed in Table 1, while walking or running on a treadmill (Fig. 2 shows the temporal variations of those six angles at six representative speeds). We set 11 different belt speeds and required participants (N=15, ages 23-31 years, all male) to walk when the belt speed was either 0.56, 0.83, 1.11, 1.39, 1.67, or 1.94 m/s and to ran when the belt speed was 2.22, 2.50, 2.78, 3.06, or 3.33 m/s.

**Table 1.** Calculated Joint angles and the definition.

| Angle | Positive value | Negative value |
| --- | --- | --- |
| Ankle | Dorsi-Flexion | Plantar-Flexion |
| Knee | Flexion | Extension |
| Hip | Flexion | Extension |

**Fig. 2.**
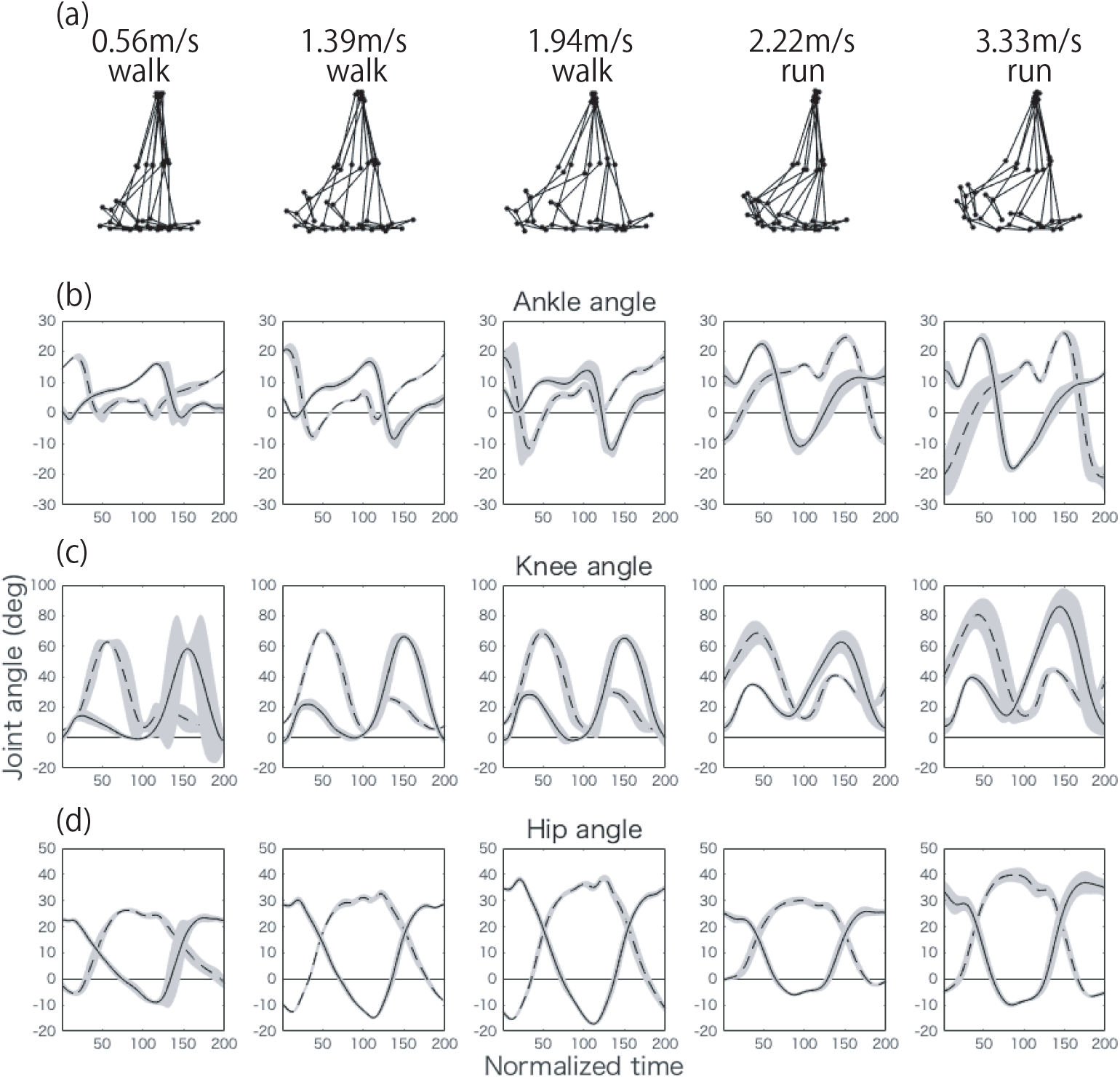
Joint angles of a representative subject at five representative speeds (0.56, 1.39, 1.94, 2.22, and 3.33 m/s). (a): Positions of the right hip, knee, and ankle every 20 time frames when normalized to 200 time frames. At time 1, the right foot took off from the ground, and it returned at time 200. (b, c, d): Non-normalized joint angles of the ankle (panel (b)), knee (panel (c)), and hip angles (panel (d)) at each speed. The dotted lines indicate those angles for the left leg, and the solid lines indicate those angles for the right leg. The lines indicate the averaged joint angle across 27 cycles, and the shaded areas indicate the standard deviation of those angles. We focused on the average joint angles in the tensor and matrix decompositions. Table 2 summarizes the meanings of the positive and negative values for each joint.

**Table 2.** Measured muscles and those abbreviation.

|  |  |  |  |
| --- | --- | --- | --- |
| TA: tibialis anterior | MG: medial gastrocnemius | LG: lateral gastrocnemius | SOL: soleus |
| PL: peroneus longus | RF: rectus: femoris | VL: vastus lateralis | VM: vastus medialis |
| BF: biceps femoris (long head) | ST: semitendinosus | AM: adductor magnus | GM: gluteus maximus |
| Gmed: gluteus medius | TFL: tensor fascia latae | RA: rectus abdominis | ES: erector spinae |

We applied the tensor decomposition to the joint angle data of all the subjects at all speeds (Fig. 3). Throughout this study, we chose the number of modules and components depending on the explained percentages of the original variance (Fig. 3a) for the comparison to the matrix decomposition. We determined the criteria to be 70% for interpretability. Although it is possible to increase the number of modules and components to explain more of the variance, it becomes challenging to interpret the full tensor. We thus determined the criteria to be 70% and the number of modules and components to be 3 for joint angle data and 6 for EMG data (see below).

**Fig. 3.**
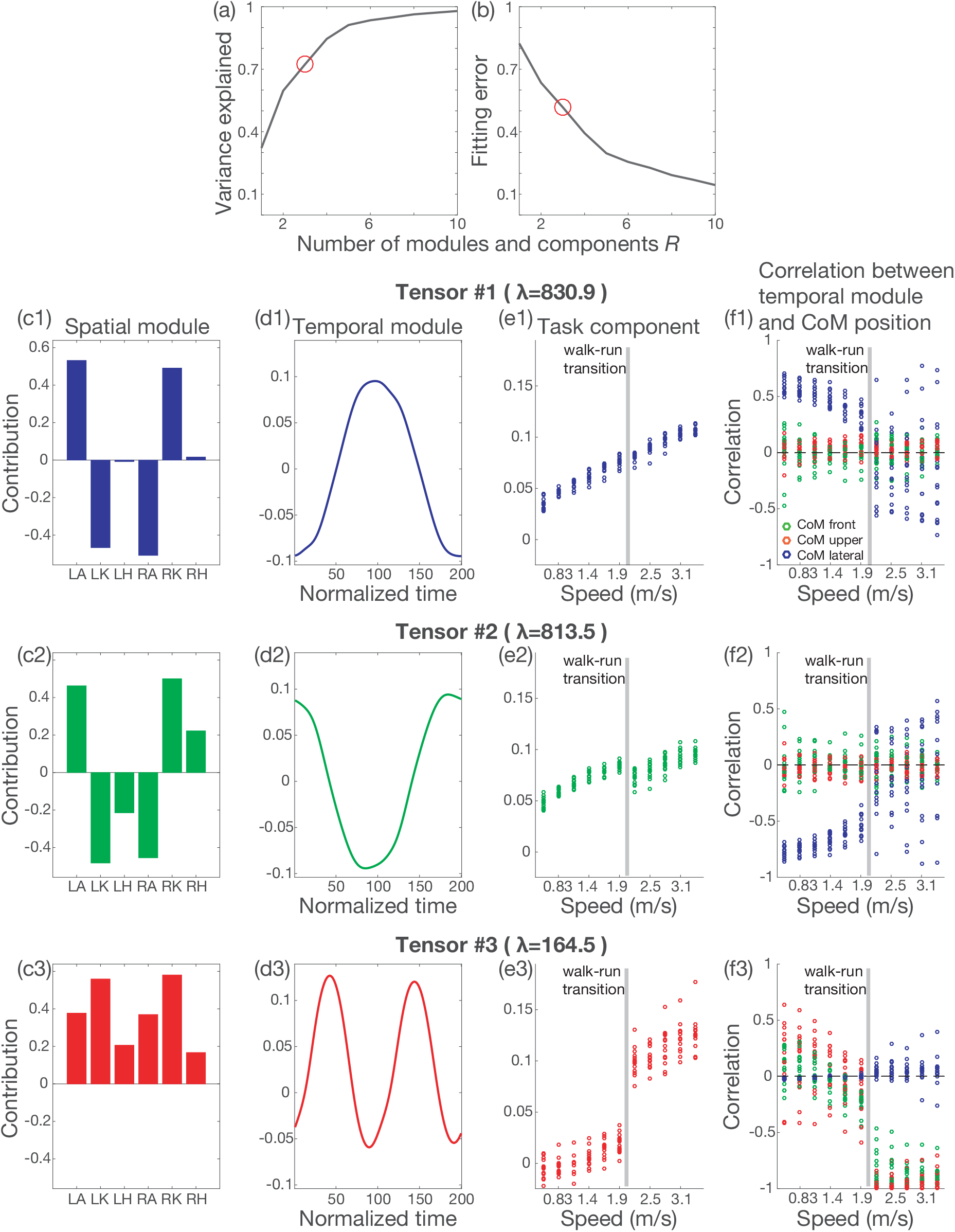
Tensor decomposition for joint angle data. (a): The relation between the number of modules and components and the variance explained by the tensor decomposition. (b): The relation between the number of modules and components and the fitting error by the tensor decomposition. (c1-c3): Extracted spatial modules. (d1-d3): Extracted temporal modules. (e1-e3): Extracted task components. (f1-f3): The correlation between the temporal module and the CoM position. λ indicates the scaling factor for each tensor. The blue, green, and red dots indicate the correlation between the temporal module and the CoM position in the lateral, frontal, and vertical axes, respectively,

Although we chose the variance explained for the comparison to the matrix decomposition, the fitting error is a more popular measure in tensor decomposition (Fig. 3b). We present the fitting error only as a reference throughout this study.

We extracted the spatial modules (Fig. 3c), the temporal modules (Fig. 3d), and the task-dependent modulations of those modules (Fig. 3e). “Task” means locomotion at the 11 different speeds for the 15 subjects. In total, we analyzed 165 tasks and obtained the respective joint angle data. We refer to each group including a spatial module, a temporal module, and a task component as a tensor hereafter (i.e., with the associated modules and components in Fig. 1a). An essential consideration in tensor decomposition, or CP decomposition, is that all tensors are unrelated. In other words, the spatial module presented in Fig. 3c1 is associated with the temporal module indicated in Fig. 3d1 and the task component presented in Fig. 3e1; however, that spatial module is not related to other spatial modules, temporal modules, or task components. Throughout this study, we indicate the associated modules and components using the same color, such as blue, green, and red, in the tensor decomposition. In the matrix decomposition (Figs. 1b and 4), the color does not always indicate associations.

**Fig. 4.**
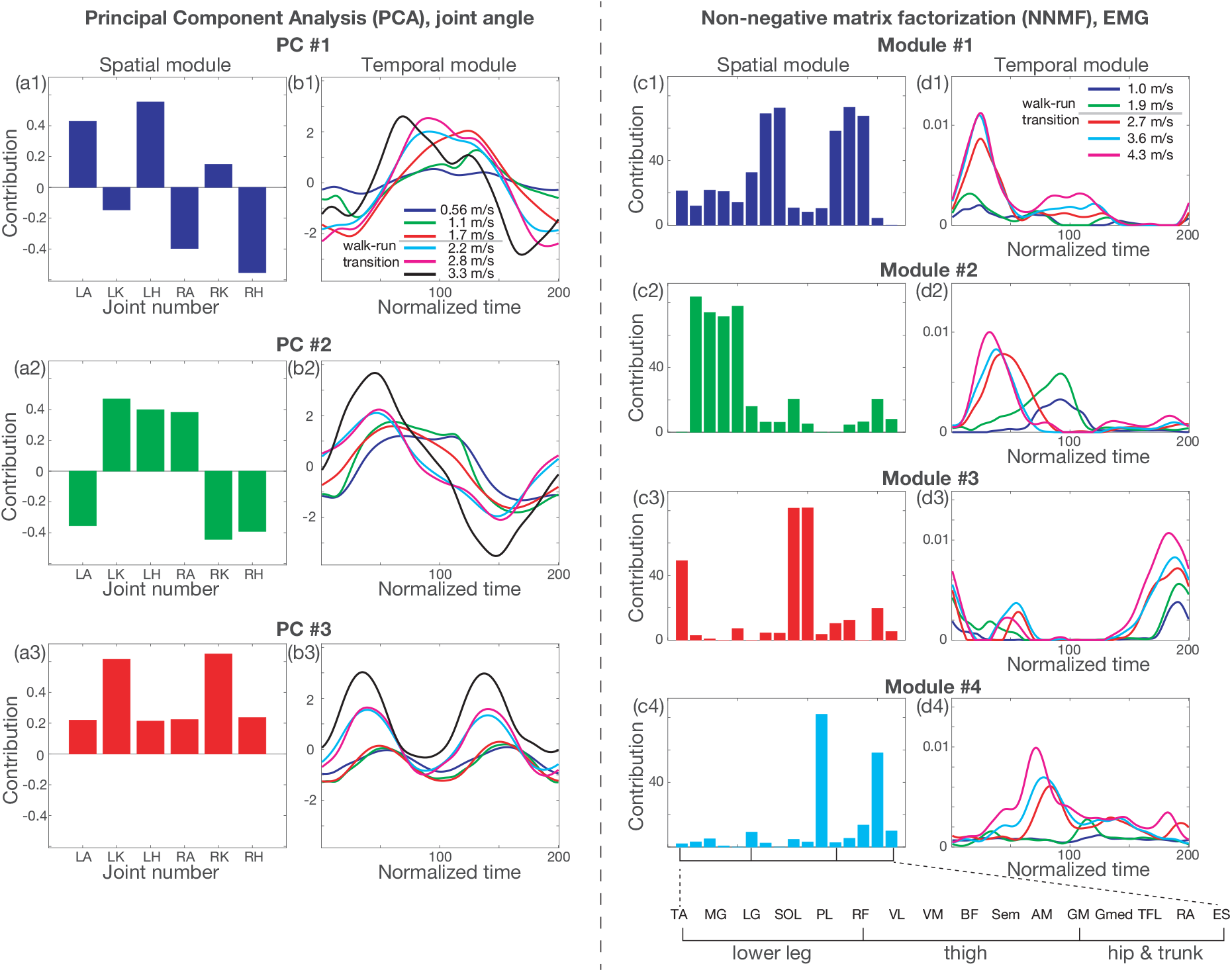
Matrix decomposition for joint angle data and EMG data. (a1-a3): Spatial modules extracted by PCA for joint angle data. (b1-b3): Temporal modules extracted by PCA for joint angle data. (c1-c4): Spatial modules extracted by non-negative matrix factorization (NNMF) for EMG data. (d1-d4): Temporal modules extracted by NNMF for EMG data.

In tensor #1 (Fig 3c1-3f1), the spatial module consisted primarily of the left ankle, left knee, right ankle, and right knee. The contributions of the left and right legs were the opposite because of the opposite sign in the spatial module. The temporal modulation was minimized when the right foot took off (at time 1) and returned (at time 200) and was maximized when the left leg was on the belt (at approximately time 100). These results indicate that the left ankle showed plantar-flexion, the left knee showed flexion, the right ankle showed dorsiflexion, and the right knee showed extension at the foot contact of the right leg. These results also indicate that the left ankle showed dorsiflexion, the left knee showed extension, the right ankle showed plantar-flexion, and the right knee showed flexion at the foot contact of the left leg. Based on the task component (Fig. 3e1), this spatiotemporal module was recruited more so at higher speeds.

In tensor #2 (Fig. 3c2-3f2), the left hip and right hip were additionally recruited in comparison to tensor #1. The temporal module showed the opposite sign, and the peak timings were slightly different from the peak timings in tensor #1. These results indicate that the temporal variation of the joint angle was opposite to that of tensor #1. Tensor #2 was recruited at a higher speed; however, the recruitment slightly and discontinuously decreased when the subject switched from walking to running.

In tensor #3, all the joints are cooperatively activated with two positive and negative peaks in the temporal module. The spatiotemporal module was recruited mainly during running (Fig. 3e3).

In summary, the tensor decomposition enabled the extraction of the spatial modules, the temporal modules, and the task-dependent modulations of those modules. For the comparison to matrix decomposition, we also applied PCA, a matrix decomposition algorithm mainly used for extracting spatiotemporal modules inherent in joint angles, to the same data (Figs. 4a and 4b). When we applied PCA as demonstrated in Fig. 1b, we obtained the spatial modules common across all speeds and subjects (Fig. 4a) and the modulated temporal modules depending on the task (Fig. 4b). One difference between the tensor decomposition and PCA is the orthogonality. In PCA, the spatial modules are orthogonal to each other; however, the orthogonality restriction originates from mathematical convenience rather than the properties of the spatial modules. The tensor decomposition yields spatiotemporal modules without this limitation. Another difference between the tensor decomposition and PCA is the quantification of the task-dependent modulations. In PCA, we somehow need to quantify the task-dependent modulations in the extracted temporal modules shown in Fig. 4b. On the other hand, the tensor decomposition can quantify the modulation across all tasks without any *a posteriori* analysis. Although a simple method to quantify the task-dependent modulation is to calculate the correlation coefficients, correlation analysis often generates pairwise similarities. The tensor decomposition can quantify the task-dependent modulation globally while considering all the tasks to be analyzed.

In addition to the qualitative interpretations of each spatiotemporal module, as mentioned above, we quantitatively evaluated the functional roles of each module by measuring the CoM position for each subject. Tensors #1 and #2 were correlated to lateral CoM positions (Figs. 3f1 and 3f2) more significantly for walking than for running, and tensor #3 was related to frontal and vertical CoM positions more often for running than for walking. As shown here, one possible way of interpreting the functional roles of each spatiotemporal module is to measure additional task-relevant information; however, the important thing is that the tensor decomposition extracted those modules and the task-dependent modulations from only motion data (i.e., joint angle data), without any additional information (e.g., CoM positions). These correlations were thus found in the *a posteriori* analysis.

### 2.3 Tensor decomposition for EMG data

Tensor decomposition can be applied to not only joint angles but also EMG data with non-negative constraints. We measured 16 muscles on the right side, listed in Table 2, as 16 subjects (ages 20-31 years, all male) walked or ran on the treadmill. The belt speed gradually increased from 0.3 to 5.0 m/s for the well-trained college runners (N=8) and 0.3 to 4.3 m/s for the non-runners (N=8). The belt speed gradually increased following a constant acceleration (0.01 m/*s*^2^). The subjects were instructed to walk or run based on their choice. As a result, the subjects switched their motion patterns from walking to running at approximately 1.9-2.3 m/s. The details of the measured data were described in our earlier study [12].

We applied the tensor decomposition and extracted six tensors (Fig. 5). Each tensor has a different functional role. In tensor #1 (Fig. 5c1-5f1), all the muscles in the lower legs, the quadriceps muscles, and all the hip muscles are activated (Fig. 5c1) upon right foot contact (Fig. 5d1). This spatiotemporal module was recruited more so at higher speeds (Fig. 5e1): the tensor decomposition enabled us to quantify the task-dependent modulation of those modules.

**Fig. 5.**
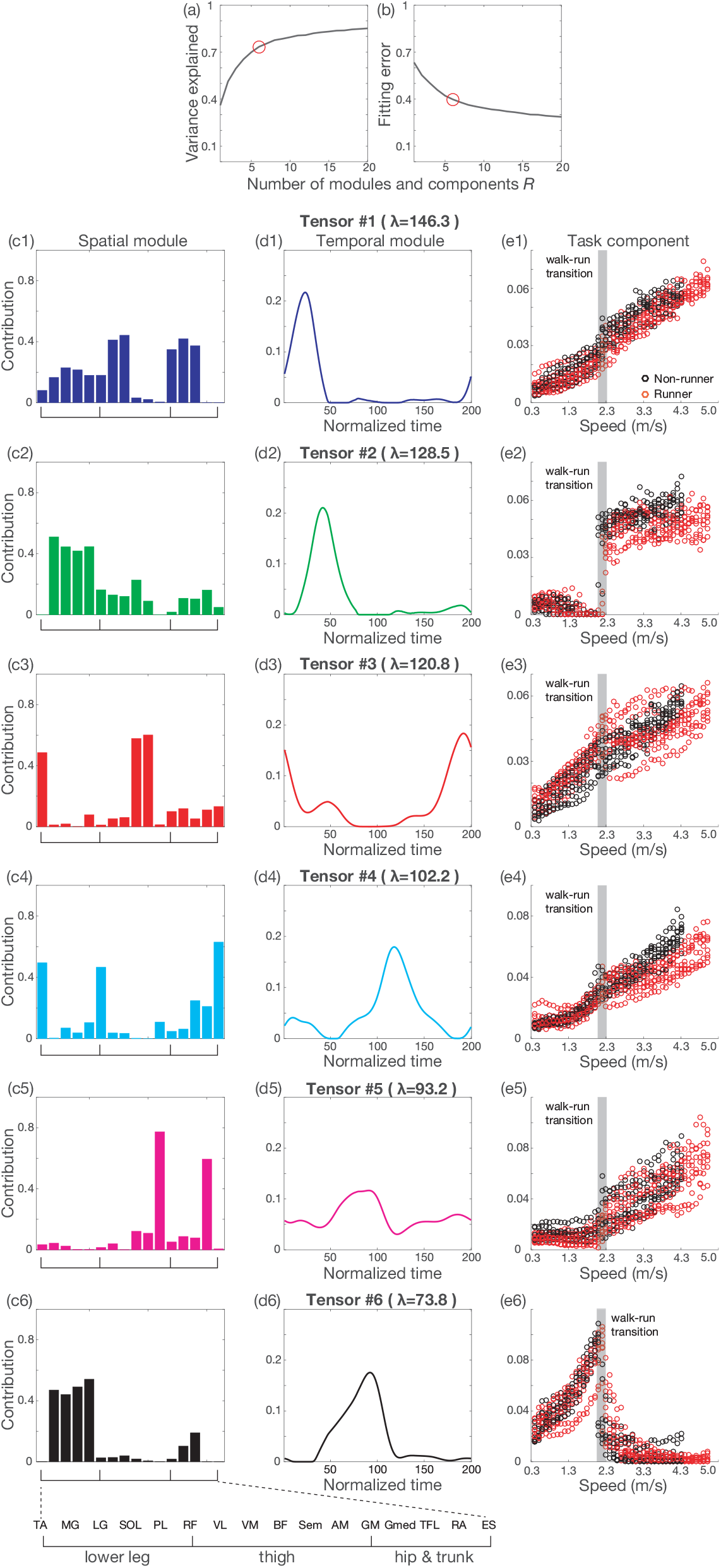
Tensor decomposition for EMG data with non-negative constraints. (a): The relation between the number of tensors and the variance explained by the tensor decomposition. (b): The relation between the number of tensors and the fitting error by the tensor decomposition. (c1-c6): Extracted spatial modules. (d1-d6): Extracted temporal modules. (e1-e6): Extracted task components. λ indicates the scaling factor for each tensor.

One difference between the tensor decomposition and NNMF is how the task-dependent modulation is quantified. In NNMF (Fig. 4c and 4d), we obtained the common spatial modules across all the tasks and the temporal modules modulated in a task-dependent manner. The NNMF requires some posterior analysis to evaluate the modulation. One popular method is to utilize the correlation coefficient. Although the method is convenient, it has limitations; the correlation coefficient can often be used to compare a pairwise relation locally ̶ in addition, NNMF in the form denoted in Fig. 1B assumed common spatial modules across all tasks. Tensor decomposition enabled us to quantify the task-dependent modulation of spatiotemporal modules without any *a posteriori* analysis.

In tensors #2 and #6, the task components showed discontinuous changes between walking and running (Figs. 5e2 and 5e6). The task components in other tensors showed continuous modulation depending on the speed. Because tensors #2 and #6 were related to either walking or running or because those tensors likely provided the neural mechanisms facilitating switching between walking and running, we further investigated the properties of tensors #2 and #6.

We then applied the tensor decomposition to the EMG data for each subject. After detecting the two tensors whose task components showed the largest and second largest changes between walking and running, we plotted the spatial modules (Fig. 6a), the temporal modules (Fig. 6b), and the task components (Fig. 6c). The task components showed discontinuous changes between walking and running. In particular, the tensor whose properties are shown in black was recruited mainly for walking, and the tensor whose properties are shown in green was recruited mainly during running (Fig. 6c). For the temporal modules, the peak timings were different between those tensors, supporting the previous hypothesis based on matrix decomposition [4, 22]: the CNS switched between walking and running by mainly controlling the peak timing of the temporal modules. In addition to those hypotheses, the tensor decomposition extracted the different spatial modules (Fig. 6a). A single asterisk denotes a significant difference with p < 0.05 and double asterisks denote a significant difference with p < 0.01 (p = 2.36 × 10^−9^ [F(15,225) = 5.44] for the interaction between the muscle factor and the other factor [walking or running], with p = 0.0200 for PL, p = 0.00847 for RF, p = 0.00987 for VL, p = 0.0318 for VM, p = 0.000218 for BF, p = 0.00990 for Sem, and p = 0.0416 for RA). The details of the statistical analysis are given in the Methods section. We found several significant differences in spatial modules that are related to walking or running. In particular, the thigh muscles are more significantly recruited for running than for walking. The tensor decomposition thus provided a new perspective on how the CNS switches between waking and running, i.e., not only through the modulation of the peak timing in the temporal modules but also through the recruitment of the proximal muscles for running.

**Fig. 6.**
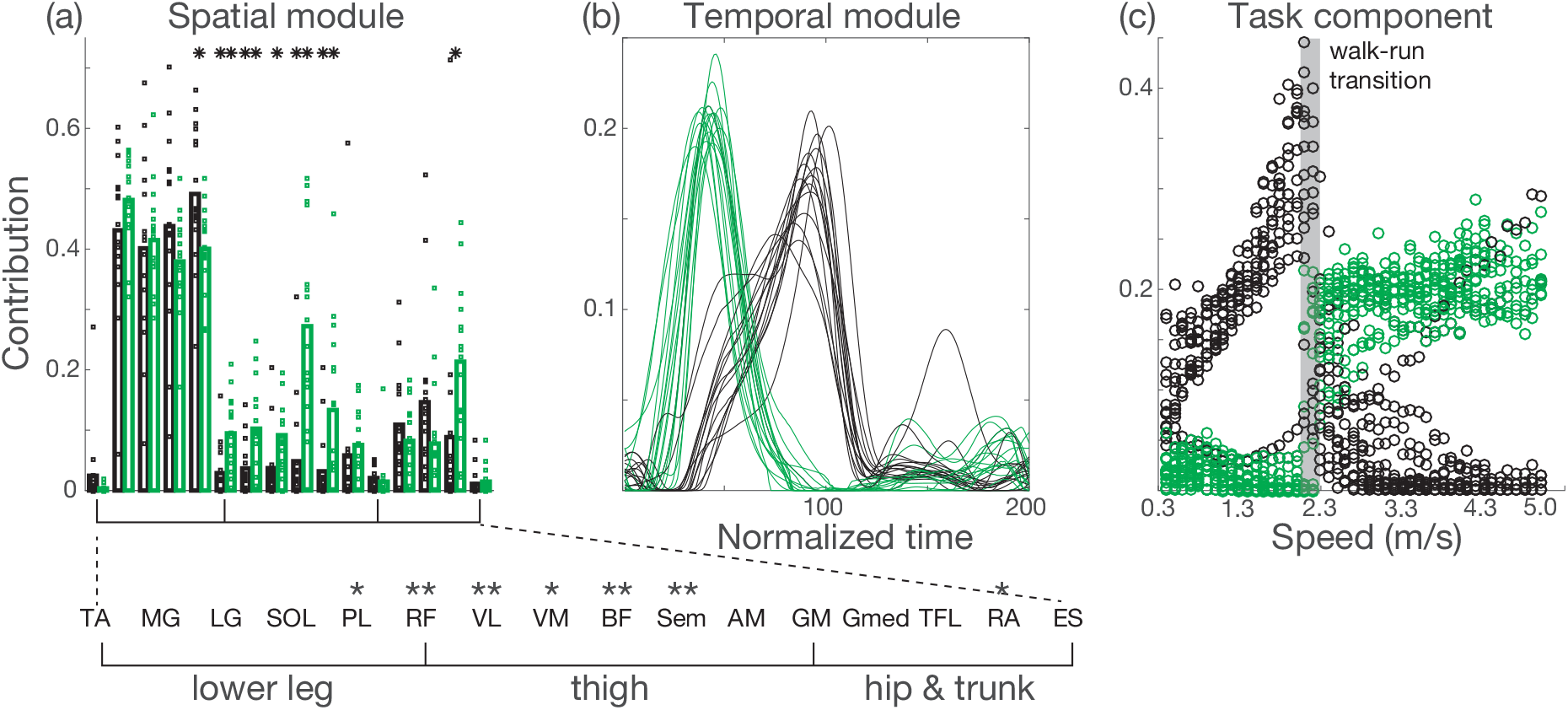
Further analysis of the tensors whose task-dependent modulations show discontinuous changes between walking and running. The tensor decomposition with a non-negative constraint was applied to EMG data for each subject. We chose the tensors whose task component showed the most and second-most substantial discontinuous changes between walking and running. We plotted the tensor that showed a more significant task component for walking than for running in black and the tensor that showed a more substantial task component for running than for walking in green. (a): Spatial modules for each subject. Each dot indicates the recruitment pattern in each spatial module and for each subject. Each bar shows the averaged values of the recruitment pattern across all subjects. Single asterisks and double asterisks associated with certain muscles indicate a significant difference in the recruitment patterns in each tensor with p < 0.05 and p < 0.01, respectively. (b): Temporal modules for each subject. (c): Task components for each subject.

## Discussion

The current study demonstrated the effectiveness of tensor decomposition in analyzing the task-dependent modulations of spatiotemporal modules on joint angle data (Fig. 3) and EMG data with non-negative constraints (Fig. 5). Matrix decomposition, such as in PCA and NNMF, is a popular method for quantifying the spatial modules, temporal modules, and task-dependent modulations of either spatial or temporal modules accompanied by *a posteriori* analysis [7–9, 11, 12]. The framework is popular not only for quantifying the task-dependent modulation but also for quantifying the individual differences [6,21,23]. As shown in this study, tensor decomposition enables the quantification of task-dependent modulations in both spatial and temporal modules (Figs. 3, 5, and 6). Although we simultaneously quantified the individual differences (e.g., runners and non-runners in Fig. 5) and task-dependent modulations, there were no significant individual differences throughout this study. Additional statistical analysis provided further information about the neural control of walking and running movements (Fig. 6). Tensor decomposition can thus be used to evaluate the task-dependent modulations and the individual differences in spatiotemporal modules in a straightforward manner.

The current study simultaneously focused on both the task-dependent modulations and individual differences of spatiotemporal modules by mixing those aspects in the “task” dimension. Because the individual differences were smaller than the speed-dependent modulations for walking and running, we clarified how spatiotemporal modules were modulated depending on the speed. When we wanted to focus on task-dependent modulations in detail, we needed to analyze tensor data under all conditions for each subject such as in the analysis of the EMG data (Fig. 6). On the other hand, when we wanted to focus on individual differences in detail, we needed to analyze tensor data under each condition for all subjects. Due to the flexibility of the tensor decomposition, we needed to carefully generate the tensor data based on the main purpose of the analysis.

The CoM position enabled us to discuss the functional roles of each tensor (Fig. 3). Because the lateral CoM position was suggested to relate to body balance in the frontal plane [24] and because its acceleration depends on the speed in a quadratic manner for elderly individuals [25], tensors #1 and #2 in Fig. 3 can play those roles mainly for walking. Other gait parameters and speed dependencies have been discussed in several studies. The speed dependence of the stride length or interval shows a U-shaped curve [26, 27]. Future work will attempt to investigate the relationship between each gait parameter and spatiotemporal module. In these analyses, however, we need to pay attention to that the tensor decomposition is an unsupervised method; the extracted spatiotemporal modules are not directly related to those gait parameters. In other words, we can extract the dimensions inherent to the joint angles or EMG data as more relevant to those gait parameters using supervised methods such as regression and classification [28]. Although it is possible to relate each spatiotemporal module to some gait parameters, we need to remember that the relation is indirect.

CP decomposition is the simplest version of tensor decomposition; it is possible to apply a more sophisticated version of tensor decomposition. A popular alternative is Tucker decomposition [15], whose variant has been applied to EMG data [17, 18]. In Tucker decomposition, the number of spatial modules, the number of temporal modules, and the number of task components can differ from each other. On the other hand, there are three free parameters (i.e., the number of spatial modules, the number of temporal modules, and the number of task components), which requires a massive computational time compared to CP decomposition. Another variant of tensor decomposition is to include a smoothness property to the tensor decomposition [29]. Because the temporal variations of the joint angle and EMG signals are smooth, the smoothness property can be used to effectively denoise the data such as in the state model in the state space model [30–33]. For the analysis of a single condition and subject, the smoothness can also be effectively applied to task components.

Because the current study focused on steady-state motion without any perturbations or unexpected changes in the environment, one possible future research direction is to apply tensor decomposition to joint angle or EMG data in response to perturbations [8,34] or trial-to-trial adaptation to perturbations [35–37] to extract response- or adaptation-related spatiotemporal modules. For extracting such error-correction-related modules, the tensor decomposition can be effectively used after being combined with regression frameworks [38]. A regression framework enables us to extract the hidden information relevant to target values [28, 39]. Because several studies have focused on adaptation [35–37, 40–43] and spatiotemporal modules [2–6, 11] separately, the relation between those concepts has only been investigated in a few studies [10, 16, 28]. Investigating the link between motor adaptation and the spatiotemporal modules via tensor decomposition represents promising future work.

## 3 Materials and Methods

### 3.1 Ethics Statement

A total of 31 healthy volunteers (ages 20-31 years, all male) participated in our experiments, which were approved by the ethics committee of the University of Tokyo and were performed following guidelines and regulations. All participants were informed of the experimental procedures following the Declaration of Helsinki, and all participants provided written informed consent before the start of the experiments.

### 3.2 Experimental setup, data acquisition, and data processing (joint angles)

A total of 15 participants participated in our experiment to measure joint angles. They performed walking at six speeds (0.56, 0.83, 1.11, 1.39, 1.67, and 1.94 m/s) and running at five speeds (2.22, 2.50, 2.78, 3.06, 3.33 m/s) on a treadmill (Bertec, Columbus, OH, USA). Under all the conditions, we measured more than 27 strides; we thus analyzed the joint angles averaged across the first 27 strides for all speeds and subjects.

The joint angles and position of the center of mass (CoM) were recorded at 100 Hz using 12 cameras (Optitrack V100: R2, NaturalPoint Inc., Corvallis, Oregon) under the marker set according to the lower body plug-in gait marker set with additional upper body markers to estimate the CoM position with less time-consuming and high accuracy [44]. The measured marker positions were low-pass filtered with a zero-lag Butterworth filter (15-Hz cut-off, 4th order) and transformed into joint angles in the sagittal plane (i.e., right and left ankle, knee, and hip angles).

Three-dimensional ground reaction force (GRF) data were recorded at 1000 Hz through force plates under each belt of the treadmill. The sampling rate was modified to 100 Hz to match the rate of the joint angles. The GRF data were low-pass filtered with a zero-lag Butterworth filter (15-Hz cut-off, 4th order). The times corresponding to foot-contact and toe-off were determined on a stride-to-stride basis from the vertical component of GRF.

Because the stride-to-stride cycle differed depending on the speed, we normalized all the cycles to 200 time frames for all speeds and subjects. Accordingly, we normalized all the joint angles. In addition, we normalized the joint angles so that the mean and standard deviation of each angle for each subject across all speeds were 0 and 1, respectively. These normalizations enabled us to compare different joint angles, speeds, and subjects fairly. In total, the joint angle data included six joint angles, 200 time frames, and 11 × 15 = 165 (i.e., the number of speeds × the number of subjects) task datasets. Throughout this study, the word “task” broadly indicates different tasks (i.e., locomotion at a different speed) for all the subjects. In this case, 165 tasks included 11 speeds (i.e., 11 types of tasks) and 15 subjects. We thus made the tensor data ***X*** ∈ **R**^6×200×165^ to apply tensor decomposition to all the data at once.

### 3.3 Experimental setup, data acquisition, and data processing (EMG)

We analyzed our earlier EMG data, the details of which can be found in [12]. A total of 16 participants participated in our experiment for measuring EMG data. A total of 8 of sixteen participants were well-trained college runners. The runners experienced higher speeds than other participants. All participants walked or ran on the same treadmill as mentioned above with linearly increasing speed (ramp speed condition, with an acceleration set to 0.01 m/*s*^2^). The speed range was adjusted to each group safely but as widely as possible (0.3-4.3 m/s in the non-runner group and 0.3-5.0 m/s in the runner group). The participants were instructed to choose to either walk or run depending on their preference under the given speed. The transition speed from walking to running for all participants ranged from 1.9 to 2.3 m/s. Because the acceleration was tiny and because the maximum speeds were considered safe for each group, the locomotive movements by all participants were always stable during the experiment.

Three-dimensional GRF data were recorded in the same manner mentioned above. The surface EMG activity was recorded from the listed 16 muscles (Table 2) on the right side of the trunk and leg. The EMG activity was recorded with a wireless EMG system (Trigno Wireless System; DELSYS, Boston, MA, USA). The EMG signals were bandpass filtered (20-450 Hz), amplified (with a 300 gain preamplifier), and sampled at 1000 Hz. The EMG data were digitally full-wave rectified and smoothed, as well as low-pass filtered with a zero-lag Butterworth filter.

Because the stride-to-stride cycle differed depending on the speed, we normalized all the cycles to 200 time frames for all speeds and subjects. Accordingly, we normalized all the EMG signals. In addition, we normalized the EMG signals so that the maximum value of each muscle for each subject across all speeds was 1. In this paper, we divided the belt speed into 0.1 m/s intervals, e.g., 0.3-0.4 m/s and 0.4-0.5 m/s. After defining the speed range, we averaged the EMG activities in each speed range. In total, the EMG data of non-runners consisted of 16 muscles, 200 time frames, and 40 × 8 tasks (i.e., 40 speed ranges and eight subjects). The EMG data of the runners consisted of 16 muscles, 200 time frames, and 47 × 8 tasks (i.e., 47 speed ranges and eight subjects). We thus made the tensor data ***X*** ∈ **R**^16×200×696^ for applying tensor decomposition to all the data. To apply the tensor decomposition to each subject, the size of the tensor data was ***X*** ∈ **R**^16×200×40^ for each non-runner and ***X*** ∈ **R**^16×200×47^ for each runner. In applying the tensor decomposition to the EMG data for each subject (Fig. 6), we set the number of tensors *R* to 6 to make the number the same in the analyses of all the subjects (Fig. 5).

### 3.4 Tensor decomposition

We relied on the tensor toolbox in MATLAB [45, 46] and used the function “cp als” (alternating least squares [15]) for analyzing the joint angles and “cp nmu” (multiplicative update, similar to NNMF [14]) for analyzing the EMG signals.

An important aspect of the tensor decomposition for joint angles (i.e., data without non-negative constraints) is that any two pairs of components can be sign reversed under the same approximated value. For example, when ***w***_*r*_ → −***w***_*r*_ and ***p***_*r*_ → −***p***_*r*_, the approximated values are invariant. If we apply the tensor decomposition to the joint angle data of two subjects separately, the sign of the spatiotemporal modules can be different even when the modules are similar for each subject except for being positive or negative. We thus apply the tensor decomposition to the joint angle data, including all the subjects, to estimate the common spatiotemporal module and task-dependent modulation of those modules for each speed and each subject.

### 3.5 Matrix decomposition

When we compared the tensor decomposition to PCA and NNMF, we relied on the MATLAB functions “pca” and “nnmf”.

To apply the matrix decomposition to the joint angle data, the size of the matrix data was ***Z*** ∈ ***R***^6×16500^, where 6 is the number of joints and 33000 is the multiplication of the number of time frames, the number of speeds, and the number of participants. To apply the NNMF to the EMG signals, the size of matrix data was ***Z*** ∈ ***R***^16×139400^, where 16 is the number of muscles and 139400 is the multiplication of the number of time frames, the number of speed ranges, and the number of participants.

### 3.6 Statistical test

To compare the task components of two representative tensors in Fig. 6, we first extracted the two tensors based on the absolute difference of the task components between walking (the data at 1.8 m/s) and running (the data at 2.3 m/s). When the task component was larger for walking than for running, all the components are written in black. When the task component was greater for running than for walking, all the components are written in green. These colors are based on the results denoted in Fig. 5. Tensors #2 (green) and #6 (black) in Fig. 5 extract task components larger for running than for walking and larger for walking than for running, respectively.

After separating the two tensors, we performed a repeated-measure ANOVA for the recruited values in the spatial modules with two factors: the type of muscle and either walking or running. After confirming the interaction between those factors, we performed Tukey’s post-hoc test.

## Acknowledgements

We thank Daichi Nozaki, Shin-ichi Furuya, Shinya Fujii, and Shota Hagio for their helpful comments.

## Notes

#### Summary of Updates

Revised mathematics. Specifically, we added scaling factor for each tensor (i.e., the contribution of each tensor to explain original data) in mathematical equations and figures.

